# CNN Model With Hilbert Curve Representation of DNA Sequence For Enhancer Prediction

**DOI:** 10.1101/552141

**Authors:** Monowar Anjum, Ibrahim Asadullah Tahmid, M. Sohel Rahman

**Affiliations:** Samsung Research, Bangladesh; Department of Computer Science and Engineering, Bangladesh University of Engineering and Technology

## Abstract

**Motivation:** Enhancers are distal cis-acting regulating regions that play a vital role in gene transcription. However, due to the inherent nature of enhancers being linearly distant from the affected gene in an irregular manner while being spatially close at the same time, systematically predicting enhancers has been a challenging task. Although several computational predictor models through both epigenetic marker analysis and sequence-based analysis have been proposed, they lack generalization capacity across different enhancer datasets and have feature dependency. On the other hand, the recent proliferation of deep learning methods has opened previously unknown avenues of approach for sequence analysis tasks which eliminates feature dependency and achieves greater generalization. Therefore, harnessing the power of deep learning based sequence analysis techniques to develop a more generalized model than the ones developed before to predict enhancer region in a DNA sequence is a topic of interest in bioinformatics.

**Results:** In this study, we develop the predictor model CHilEnPred that has been trained with the visual representation of the DNA sequences with Hilbert Curve. We report our computational prediction result on FANTOM5 dataset where CHilEnPred achieves an accuracy of 94.97% and AUC of 0.987 on test data.

**Availability:** Our CHilEnPred model can be freely accessed at https://github.com/iatahmid/chilenpred

## 1. Introduction

An Enhancer is a short region of DNA that can be bound by proteins to increase the likelihood that transcription of a particular gene will occur (12)(13). These proteins are usually referred to as transcription factors. In higher eukaryotes, enhancers activate gene transcription by recruiting transcription factors and their complexes and in doing so, they contribute to vital biological processes including development and differentiation (14)(15), maintenance of cell identity (16), response to stimuli (18)(17), and interactions with target genes through promoter-enhancer looping (19)(20)(21). Also, genetic disruption in enhancers has been found to be closely associated with cancers (22).

Although enhancers play a vital role in various genetic phenomena, a promising way of identifying novel enhancers is yet to be found. There are several reasons for this (13). First, the number of enhancer sequences are very small as compared to the size of the human genome. Second, enhancers are distal cis-acting DNA regions, meaning, they can be located far away from the gene, upstream or downstream from the gene they regulate. They do not necessarily act on the respective closest promoter but can bypass neighboring genes to regulate genes located more distantly along a chromosome. Third, in contrast to the well-defined sequence code of protein-coding genes, there is no general sequence code of enhancers. Thus, identifying enhancer computationally has become a challenging task.

The versatility and the complexity of the datasets that are available through recent experimental procedures do not make this task any easier. For instance, Chromatin Immuno-Precipitation followed by massive Sequencing (ChIP-Seq) sheds light on chromatin accessibility in different organisms, tissues and under different conditions. On the other hand, Cap Analysis of Gene Expression (CAGE) estimates the quantity of 5′ ends of messenger RNA in a cell. Projects, such as the ENCODE (32) and the NIH Epigenome Roadmap (33), released libraries of histone modification marks in the human genome, whereas the FANTOM5 project (34) released CAGE-based transcription start sites (TSSs) in different cell types and tissues and enabled for the comprehensive identification of functional regulatory elements. Thus, developing a computational model to derive relevant information from these datasets has become essential. The problem of identifying enhancers can be defined as follows: given a DNA sequence, predict if it can function as an enhancer (23). Numerous computational methods have been proposed for improving the enhancer prediction.

One of the features that pioneered the computational approaches for enhancer prediction is based on evolutionary conservation (2). Visel et. al (3) argue that human regulatory elements show low conservation among different species. As a result, they can not be characterized with confidence based on enhancer regions in other mammals. The second category of computational methods relies on more sophisticated algorithms that associate enhancers and promoters with certain types of histone modification marks and transcription regulators (18). However, since the types of histone modifications and regulators that identify enhancers differ significantly, they lack the generalization of the prediction model. The third category of methods approach the enhancer detection problem as a binary classification task by discriminating enhancer regions from non-enhancer regions using supervised machine learning techniques, such as support vector machines (SVMs) (24)(23), artificial neural networks (ANNs) (25), decision trees (DTs) (26), random forests (RFs) (27), and, more recently, deep learning (29). On the other hand, the unsupervised learning approaches like ChromHMM(28) offer genomic segmentation and characterization based on a Hidden Markov Model (HMM) whereas Segway(30) presents a dynamic Bayesian network. Solely depending on the sequence-based analysis, BiRen (35) offers a hybrid network of Convolutional Neural Network (CNN) and Bidirectional Recurrent Neural Network. DEEP (10) presents an ensemble technique to train classifiers with unbalanced classes.

In our background studies on enhancer prediction, we have not found any methodology that explores the idea of representing the DNA sequence in a different way. However, some studies have been found to visually represent a DNA sequence with a discussion on their impact on classification task. Anders (11) first proposed the idea of visualizing the genomics data with Hilbert Curve, one form of a space-filling curve. Later, Yin et. al. (1) perfected on that idea and showed that such visual representation helps to develop a CNN that performs significantly well on classification tasks in bioinformatics.

In this article, we have developed a CNN model named CHilEnPred that integrates the representational power of the Hilbert Curve with the spacial hierarchical information extraction power of CNN. CHilEnPred has been trained with FANTOM5 Human Enhancer dataset which applies a distinct bidirectional CAGE pattern to identify enhancers in human tissue and cells. We demonstrate that our model illustrates superior prediction accuracy relative to the state-of-the-art methods based on sequence characteristics. Since our model does not require any feature engineering and depends solely on the image representation of the DNA sequence, it has the power to generalize to other species as well. Our CHilEnPred model has opened a new avenue of exploration in the study of predicting distal cis-acting DNA regions like enhancers. We believe it will provide the researchers with a broader understanding of the characteristics of enhancer sequences as well.

## 2. Methods

### 2.1 Hilbert Curve

Our initial approach was to convert the DNA sequence in an image which had the height of the length of the sequence and the width would be 4 where each column would correspond to a specific base {’A’,’T’,’C’,’G’}. We tried various combinations of CNN and RNN to predict enhancers. However, none of them showed a promising result or significant improvement over existing results. Feature engineering of the enhancer sequences was also ineffective for this purpose. After these efforts, we started to rethink our approach from the beginning again. The unique relational formation of the enhancers and the regulated sequences encouraged us to do so. In Eukaryotic cells, the structure of the chromatin complex of DNA is folded in a way that functionally mimics the super-coiled state characteristic of prokaryotic DNA, so although the enhancer region in the DNA sequence may be far from the gene in a linear way, it is spatially close to the promoter and gene (31). We need a visual representation that does not rely on the linear representation of the sequence, and perfectly captures such distal characteristic of the enhancers. We hypothesize that it can be achieved by a space-filling curve which folds a 1-D linear sequence in a 2-D image. So, as seen from Figure 1, although the enhancers are found far away from the regulated gene, in a folded 2-D space-filling curve, they are closer.

**Fig. 1:**
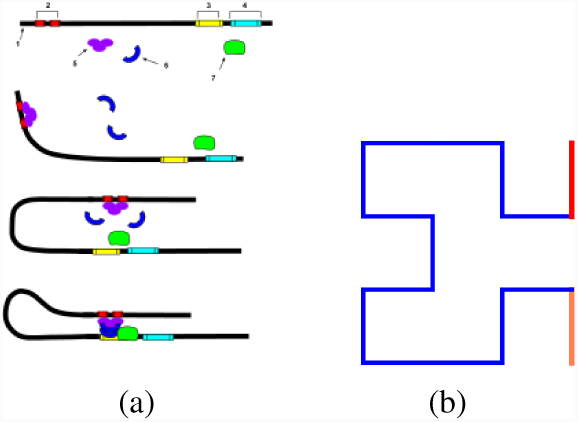
(a) Enhancers are linearly far away from the gene, but closer in a folded structure. (b) Two nodes having a linear distance of 15 units comes next to each other in a Hilbert Curve of order 2.

### 2.2 Image Representation of the DNA sequence

Each of the enhancer sequence samples we have used in our experiment has a sequence length of 401. We transformed these sequences into images by following three distinct steps namely, *k*-mer construction, one-hot vectorization, and image construction based on the Hilbert Curve which are described in details below.

DNA sequences are composed of nucleotides; however, these nucleotides do not have significant meaning when treated individually. Therefore, it is common in molecular biology problems to treat DNA sequences as a sequential collection of *k*-mers. Given that the alphabet of a DNA sequence consists of 4 letters i.e. {A,T,C,G}, the number of possible *k*-mers for any given *k* is 4^*k*^. This representation of a DNA sequences as a *k*-mer list is more suitable for statistical analysis as it is a common technique in text mining and other natural language processing tasks. For example a DNA sequence of “ACCTATAT” can be represented as a list of 3-mers: {“ACC”,“CCT”,“CTA”,“TAT”,“ATA”,“TAT”}. In the first step, we took each of our DNA sequence of length 401 and transformed them into a list of *k*-mers. We did some primary experiments for determining the appropriate value of *k*. During these experiments, we noticed that higher values of *k* were more prone to overfitting in our specific case as our dataset was significantly small. Therefore, we decided to use *k* = 1 for our specific problem.

**Fig. 2:**
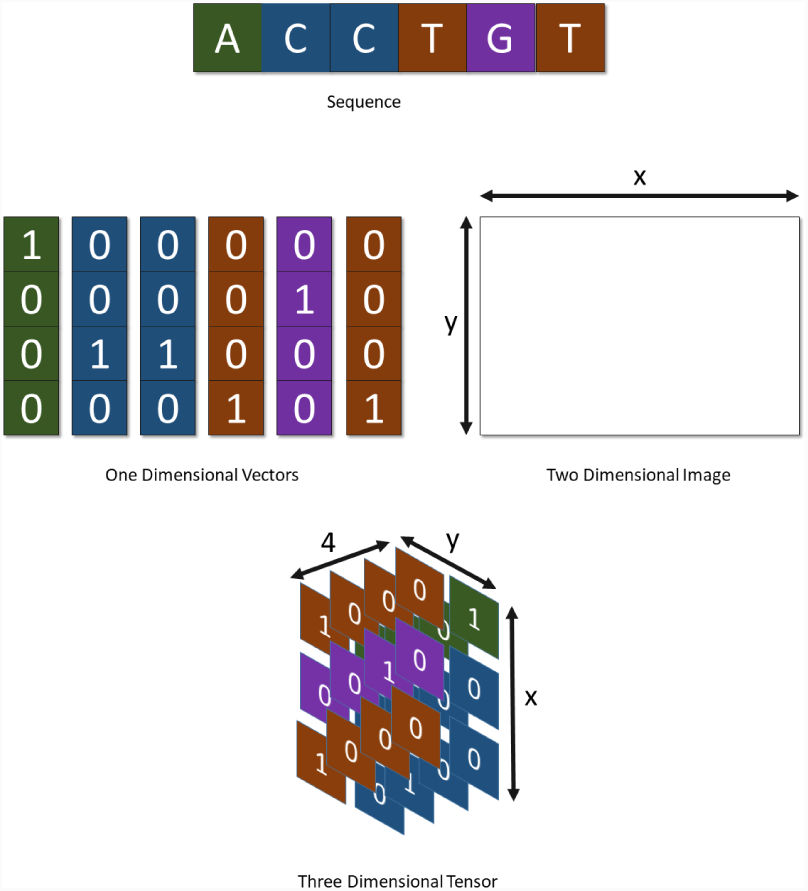
Image Representation Of DNA Sequence.

After completion of the first step, we had a discretized representation of the sequence. In the second step, we transformed that discretized representation to the mathematical form by using vectorization. In natural language processing, it is common to transform words into the mathematical form by using either word embedding, such as, word2vec(36), GloVe(37) or by using one-hot vectors. However, word embedding is not appropriate in our case. Word embedding tend to match similar meaning words closer in vector space. Since DNA sequence *k*-mers are not expected to have any meaningful co-relation with each other, we decided not to use word embedding to mathematically represent the *k*-mer list. We used one hot vector instead. One hot vector is a vector in which each position corresponds to a single *k*-mer. Therefore, the length of the vector is 4^*k*^. For example, if *k* = 1 then the one hot vector will have length 4. Here, ‘A’ will correspond to [1, 0, 0, 0], ‘G’ will correspond to [0, 1, 0, 0], ‘C’ will correspond to [0, 0, 1, 0] and ‘T’ will correspond to [0, 0, 0, 1].

After completion of the second step, we had a list of one hot vectors which represented the DNA sequence. In our third step, we transformed these one hot vectors into an image representation by using Hilbert Curve which is explained in details here. To transform the list of one-hot vectors to an image we have to find a mapping to each of the one-hot vectors to a specific pixel of the image. Since we are mapping one-dimensional vectors to a two-dimensional image, the representation becomes a 3-dimensional tensor. However, unlike RGB images that have 3 channels, this image representation has 4 channels (as the length of the one hot vectors is 4^1^ = 4 for *k* = 1).

There are various space-filling curves available which maps 1-dimensional sequences to a two-dimensional plane. However, given the task at hand one specific curve stands out which is known as the Hilbert curve. Yin et al has shown in (1) that Hilbert Curve shows better performance over other space-filling curves when it comes to the task of creating image representation from one hot encoded DNA sequence. The construction of the Hilbert curve follows a recursive pattern. In the first iteration, a Hilbert curve is drawn and is divided into 4 parts. Each part is assigned to one of the four quadrants of a square. Each quadrant subsequently gets divided into four part and each part of the initial Hilbert curve gets subdivided accordingly. Therefore, each subdivision of the square holds 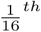 of the curve. The process can be farther continued. For example, after the next iteration, each subdivision of the square will hold 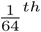 of the curve. Since on each iteration, the number of parts in the square quadruples, it can be inferred that the square holds 2^*n*^ * 2^*n*^ parts. Each part holds a corresponding part of the Hilbert curve. Since DNA sequences in our dataset are of length 401, we chose the value of *n* = 5, since choosing *n* = 4 yields an image of 256 pixels which is not suitable for holding information of the whole sequence. However, in our case, a large portion of the image remain unused as the image representation has 1024 pixels but we use only 401 of them. We did not crop the image to discard the irrelevant parts since the neural network eventually learns to discard those irrelevant pixels anyway.

**Fig. 3:**
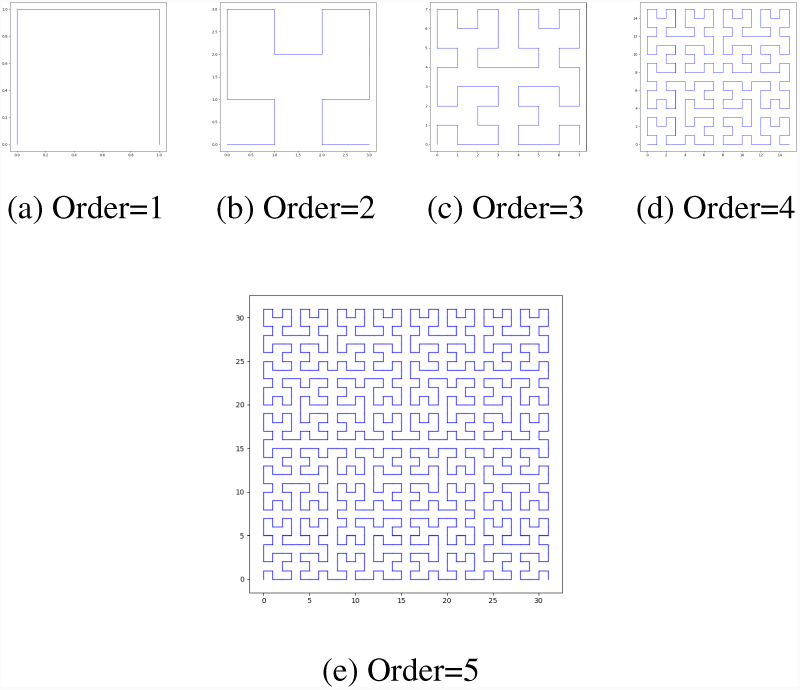
Hilbert Curves for Different Orders.

### 2.3 Network Architecture

For image recognition and representation learning tasks, Convolutional Neural Networks are widely used because of their multi-layer hierarchical feature extraction capability. The network is composed of multiple layers of parameterized kernel convolutions on the image which work to extract increasingly abstract features from the image as the network gets deeper. Upon optimizing these parameters, the CNN can be used as a universal feature extractor for images. Our network architecture is based on CNN with Fully connected layers on top of it. It is described in details in the coming sections.

We are working on images with the dimension of 32 × 32 × 4. In most CNN based models square window sizes of 3,4,5 used. In our model, the first convolution layer uses a square window of size 3. In the subsequent convolution layers, we have used the same window size. We have used rectified linear units (ReLU) as the activation function. We have also experimented with other activation functions. However, rectified linear units performed the best. For pooling layers, we have tried both average-pooling and max-pooling. However, max-pooling has shown a slight edge over the average-pooling. Therefore, in our final model, we have used max-pooling layers. These convolution layers were followed by fully connected layers with 50% dropout added. For loss function, we used binary cross-entropy.

### 2.4 Experiments

For evaluating the performance of our proposed approach we used the FANTOM5 human enhancer dataset which is publicly available (38)(39). The dataset contains 32693 human enhancer samples. Each of the samples is of length 401. Each sample in this dataset is labeled as a “positive” sample. For the negative sample, we used another dataset of random DNA strings. Negative Dataset consisted of 36800 samples.

We randomly choose approximately 75% of the dataset for training purposes. The other 20% are used for the validation test and 5% is used as the test set. We choose RMSprop optimizer for training the model. The next hyperparameter we choose is the learning rate. We initially set it to 10^-6^ which results in poor performance. We updated the value to 2 × 10^-5^ which gave the optimal training performance. We have experimented with various batch sizes and found that batch size of 256 gives the best performance in terms of generalization and reducing overfitting.

We experimented with the number of epochs and found that setting epoch number of 250 is suitable for our purpose since that is when loss becomes lowest. The performance metric of our final model was the accuracy of the model.

## 3. Results

The results of our model, CHilEnPred, for enhancer classification is very promising. Our goal was to predict enhancer from DNA sequences. We have achieved 94.97% accuracy on the test data which was separated from the training and validation data at the very beginning. AUROC of the model is 0.987. Our model shows significant performance improvement over other models which can be seen from table 1. Our proposed method conclusively outperforms both Bi-Ren and DEEP-VISTA in terms of accuracy of prediction and overall area under the curve.

**Table 1.** Network Architecture for Enhancer Classification in FANTOM5 Dataset

| Layer | Description | Output |
| --- | --- | --- |
| Input Layer |  | 32,32,4 |
| Conv2D | $3 \times 3$ Conv | 30,30,64 |
| Maxpooling2D | $2 \times 2$ | 15,15,64 |
| Conv2D | $3 \times 3$ Conv | 13,13,128 |
| Maxpooling2D | $2 \times 2$ | 6,6,128 |
| Conv2D | $3 \times 3$ Conv | 4,4,256 |
| Maxpooling2D | $2 \times 2$ | 2,2,256 |
| Flatten |  | 1024 |
| Dropout | Random 50% activation of neurons | 512 |
| Dense Layer | Fully connected layer | 512 |
| Dense Layer | Sigmoid activated output layer | 1 |

**Table 2.** Performance comparison with existing methods on accuracy and AUC

| Algorithm | Accuracy | AUC |
| --- | --- | --- |
| CHilEnPred | 94.97% | 0.987 |
| BiRen | N/A | 0.956 |
| DEEP-VISTA | 90.2% | 0.894 |
| Lee's SVM | N/A | 0.662 |

**Fig. 4:**
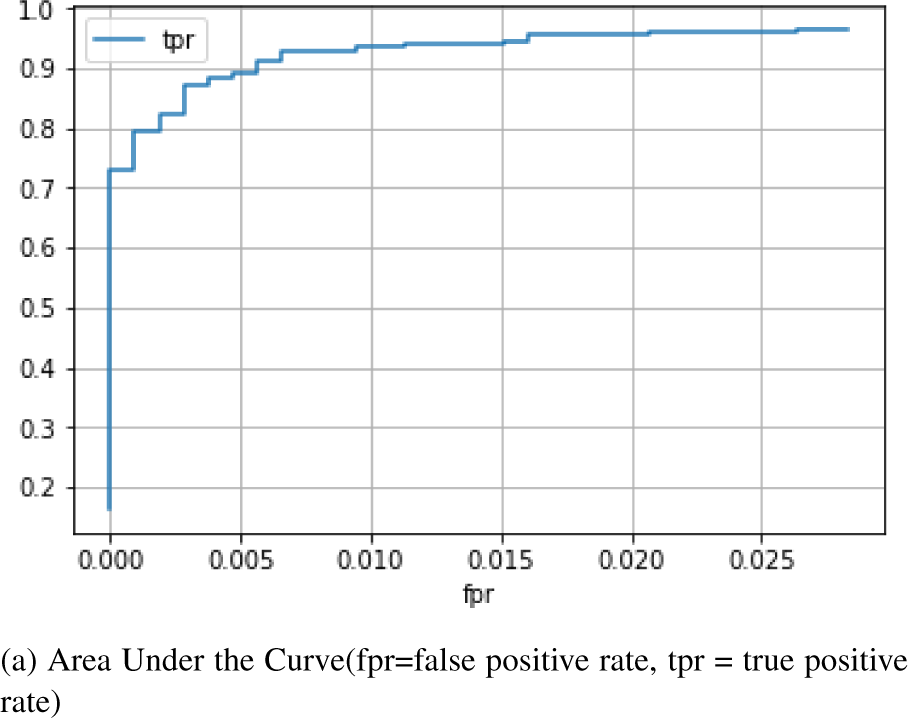
Model performance.

**Fig. 5:**
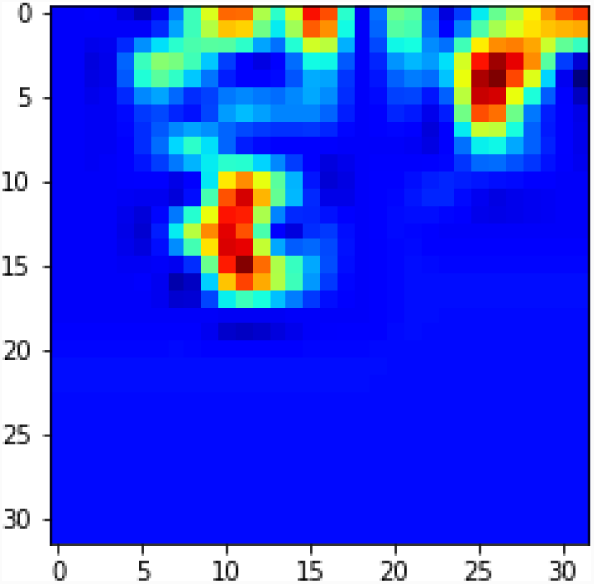
Sample Heatmap of an Enhancer Sample.

In order to have a deeper understanding of the network’s learned features, we drew the heatmaps for each enhancer. These heatmaps show a significant increase in hue in some areas other than the whole image. We find the equivalent DNA sequences from those activated pixels. Some samples of these sequences along with their occurrence count across the dataset are listed in table 3. We hypothesize that these sequences hold significant meaning in identifying enhancers. The proof for that is left for further research.

**Table 3.** Activated Feature Sequences of the model Found From The Heatmaps

| Features | Number of occurrence |
| --- | --- |
| AAA | 5012 |
| TTT | 4981 |
| CCC | 2613 |
| GGG | 2387 |
| TCT | 2136 |
| CTC | 2068 |
| CCT | 1931 |
| TTC | 1832 |
| CTT | 1793 |
| AGA | 1767 |
| GAG | 1759 |

## 4. Discussion

In this paper, we have proposed a Deep learning model which can predict enhancers from only DNA sequence information. Our method of prediction does not require any feature extraction from DNA sequences in the pre-processing phase. In most previous works, the feature extraction part was very context specific and therefore was harder to generalize. However, our proposed method is a complete end to end model where the input is an image representation of the enhancer using the Hilbert Curve and the output is the enhancer prediction. Therefore, it has more generalization capacity than the other shallow and deep learning models for enhancer prediction.

The robustness of our proposed method can be attributed to the representation of the DNA sequence. To elaborate, in the case of an enhancer, there is sizable variance in the temporal domain which means that the base pair distance between the regulating site (enhancer) and the regulated region is not fixed. Therefore, when RNN or similar models tend to capture the temporal properties of the enhancer DNA sequences they generally fall short. However, in our case, due to the inherent nature of representation of the Hilbert curve, this particular information of variance in base pair distance was better represented and therefore, resulted in a greater prediction accuracy.

## 5. Conclusion

We proposed a generalized deep learning method which can predict enhancer from DNA sequence input. Our method takes advantage of better representation by the Hilbert curve and uses CNN to predict enhancer. It shows significant performance improvement over existing methods as demonstrated by experimental results over FANTOM5 human enhancer dataset. However, there is still room for farther improvements which we would like to address in our future work. For future work, a probable avenue would be using higher dimensional images with sparse data density as data points and using specialized deep learning model on them to build a more generalized model. Furthermore, a more generalized version of our model can be achieved by incorporating datasets from species other than humans.

## Supporting information

Common Sub-sequence List

## Acknowledgements

Supported by a Titan GPU provided by university academic program.

