## Supplementary material for "CNN Model With Hilbert Curve Representation of DNA Sequence For Enhancer Prediction": Common Sub-sequence List

### Common Subsequence Found From Heatmaps

| substring | occurance |
|---|---|
| AAA | 5012 |
| TTT | 4981 |
| CCC | 2613 |
| AAAA | 2533 |
| GGG | 2387 |
| TTTT | 2289 |
| TCT | 2136 |
| CTC | 2068 |
| CCT | 1931 |
| TTC | 1832 |
| CTT | 1793 |
| AGA | 1767 |
| GAG | 1759 |
| TCC | 1752 |
| AGG | 1695 |
| CTG | 1545 |
| AAG | 1487 |
| GGA | 1479 |
| AAAAA | 1474 |
| GAA | 1472 |
| TGT | 1424 |
| GCC | 1383 |
| CAG | 1360 |
| GGC | 1312 |
| ATT | 1303 |
| ACA | 1240 |
| AAT | 1237 |
| TTTTT | 1235 |
| TGG | 1233 |
| TTA | 1195 |
| CCA | 1177 |
| GTG | 1162 |
| CAC | 1137 |
| TCA | 1097 |
| TAA | 1062 |
| TTG | 1043 |
| TGA | 1001 |
| CAA | 1001 |
| TAT | 959 |
| GTT | 937 |
| ATA | 935 |

| | |
|---|---|
| AAAAAA | 913 |
| AGT | 850 |
| GCT | 850 |
| ACT | 832 |
| TGC | 811 |
| AAC | 798 |
| GGT | 796 |
| CAT | 770 |
| GCA | 761 |
| ACC | 758 |
| AGC | 746 |
| TTTTTT | 746 |
| GTC | 745 |
| ATG | 730 |
| CCCC | 715 |
| CGG | 703 |
| CCG | 697 |
| GGGG | 661 |
| GAC | 644 |
| AAAAAAA | 601 |
| TTCT | 599 |
| CGC | 594 |
| GCG | 591 |
| CTCC | 585 |
| ATC | 567 |
| TTTC | 564 |
| TCTT | 549 |
| CTTT | 546 |
| CCTC | 535 |
| CTCT | 519 |
| GAT | 512 |
| GAGG | 510 |
| GGAG | 510 |
| CTA | 506 |
| TTTTTTT | 504 |
| CCCT | 503 |
| AAAG | 491 |
| TAG | 491 |
| TCTC | 478 |
| GAAA | 469 |
| AGAA | 468 |
| TAC | 467 |
| AAGA | 459 |

| | |
|---|---|
| TCCC | 458 |
| TCCT | 454 |
| AGGG | 453 |
| AGAG | 431 |
| AAAT | 422 |
| ATTT | 413 |
| CTTC | 405 |
| TTTA | 404 |
| AGGA | 403 |
| GCCC | 402 |
| GGGA | 399 |
| GGGC | 397 |
| GAGA | 394 |
| TTCC | 385 |
| AAAAAAA | 383 |
| TAAA | 377 |
| GTA | 372 |
| CCTT | 369 |
| TTTG | 359 |
| TGGG | 350 |
| CAAA | 346 |
| TGTT | 344 |
| CCCA | 343 |
| GGAA | 337 |
| TTTTTTTT | 334 |
| CCTG | 325 |
| TATT | 323 |
| TGTG | 320 |
| GTTT | 319 |
| GAAG | 317 |
| TTAA | 312 |
| GGCC | 309 |
| CTGG | 304 |
| TCTG | 303 |
| AATA | 301 |
| AAGG | 300 |
| CAGG | 299 |
| AACA | 295 |
| TTAT | 294 |
| TTGT | 289 |
| CCAC | 288 |
| GTGG | 274 |
| CTGC | 269 |

| | |
|---|---|
| CTGT | 267 |
| CAGA | 267 |
| AAAC | 266 |
| CACA | 263 |
| GCAG | 263 |
| CTCA | 262 |
| GTGT | 258 |
| ACAA | 257 |
| CCCG | 252 |
| AATT | 246 |
| AAAAAAA | 246 |
| ATAA | 246 |
| CGGG | 245 |
| CCAG | 244 |
| CCGC | 243 |
| ACAG | 241 |
| TTCA | 238 |
| GGGT | 236 |
| GGTG | 235 |
| CACC | 234 |
| GGCG | 232 |
| TGAG | 230 |
| GCGG | 229 |
| TTTTTTTT | 229 |
| ACCC | 226 |
| GCCT | 224 |
| CGT | 223 |
| TGCC | 222 |
| GCTG | 219 |
| GGCT | 216 |
| ATAT | 213 |
| AGGC | 210 |
| ACAC | 210 |
| GTCT | 209 |
| CAGC | 209 |
| TTTCT | 208 |
| CACT | 208 |
| TCG | 207 |
| ACG | 207 |
| TCTTT | 206 |
| TCAG | 205 |
| TTCTT | 204 |
| CTGA | 204 |

| | |
|---|---|
| TGAA | 202 |
| CGCC | 202 |
| CATT | 202 |
| TGTC | 197 |
| AGCC | 197 |
| TCAC | 196 |
| GGCA | 195 |
| AGTG | 191 |
| ATTA | 190 |
| TCAT | 189 |
| GGAGG | 187 |
| CCTCC | 186 |
| TCCA | 186 |
| GACA | 180 |
| AATG | 177 |
| AAGT | 174 |
| AAAGA | 174 |
| TCAA | 174 |
| ACTT | 173 |
| CTTTT | 170 |
| CTCCC | 170 |
| TTTTC | 169 |
| GAGT | 169 |
| CCGG | 169 |
| TAAT | 168 |
| TATA | 168 |
| TGGC | 168 |
| ACTC | 167 |
| CCCCC | 165 |
| GTGA | 165 |
| CGA | 165 |
| GAAAA | 164 |
| TGGA | 164 |
| CCCTC | 163 |
| AGAAA | 163 |
| GGGAG | 162 |
| AGCA | 162 |
| ATCT | 160 |
| AAAAG | 160 |
| TGCT | 159 |
| AGAT | 159 |
| GCCG | 158 |
| TGGT | 157 |

| | |
|---|---|
| GCTC | 157 |
| TCTCT | 156 |
| TTGA | 156 |
| GAGC | 156 |
| GAGGG | 156 |
| ATGA | 155 |
| GCCA | 155 |
| AAGAA | 155 |
| ATTC | 154 |
| TTTTTTTT | 154 |
| GGGGG | 153 |
| ATGT | 153 |
| AGAC | 152 |
| CTCTC | 148 |
| CTTA | 147 |
| AAAAAAA | 147 |
| GGTT | 146 |
| ACAT | 146 |
| TGTTT | 145 |
| CTTG | 143 |
| CAGT | 143 |
| ACCT | 142 |
| CAAG | 141 |
| TCTA | 141 |
| GGGGC | 141 |
| GTCC | 140 |
| TTGG | 140 |
| CCAA | 140 |
| ACTG | 139 |
| AAAAT | 139 |
| TTTTA | 139 |
| TAAAA | 138 |
| AGTT | 137 |
| TTAC | 137 |
| GCTT | 137 |
| AGTC | 137 |
| CGGC | 136 |
| AGAGA | 134 |
| GCCCC | 134 |
| CTCCT | 134 |
| ATTG | 133 |
| GTCA | 133 |
| AGGT | 131 |

| | |
|---|---|
| TCCCC | 131 |
| CAAT | 131 |
| CCCCT | 131 |
| CTTCC | 130 |
| AGGAG | 130 |
| GAAT | 130 |
| ATTTT | 128 |
| TCCTT | 128 |
| CATC | 127 |
| TACA | 125 |
| TTCCT | 125 |
| AACT | 124 |
| CCAT | 124 |
| AAACA | 123 |
| AATC | 122 |
| GCGC | 122 |
| GACT | 122 |
| GTTC | 122 |
| AGGGG | 120 |
| TATTT | 120 |
| CTTCT | 119 |
| CAAAA | 119 |
| AAATA | 119 |
| GGAC | 119 |
| TTTAT | 118 |
| AGGAA | 118 |
| AAGC | 118 |
| TAAG | 117 |
| TGAC | 117 |
| GAGGA | 117 |
| GATT | 116 |
| TGAT | 115 |
| TTATT | 115 |
| TAGA | 115 |
| GAGAG | 114 |
| CTAA | 114 |
| GATG | 113 |
| CCTCT | 113 |
| CTTTC | 113 |
| TTAAA | 113 |
| TCCCT | 113 |
| TACT | 112 |
| TTTTG | 112 |

| | |
|---|---|
| ATGG | 112 |
| TTTGT | 112 |
| TTGC | 109 |
| TGTGT | 109 |
| TGTA | 109 |
| GGCGG | 109 |
| ACAAA | 108 |
| TGGGG | 108 |
| GCAA | 108 |
| TGCA | 107 |
| CCCCA | 107 |
| TTAG | 106 |
| TCTTC | 106 |
| GTTG | 106 |
| CATA | 106 |
| GGAAG | 106 |
| TATG | 106 |
| TTGTT | 105 |
| AGCT | 105 |
| CCCCG | 105 |
| CATG | 105 |
| ATCA | 105 |
| CGCG | 104 |
| GGTC | 103 |
| CTCTT | 102 |
| CCCGC | 102 |
| GTTA | 101 |
| CCTTC | 101 |
| AGAGG | 101 |
| TTTTTTTT[illegible] | 101 |
| ATCC | 100 |
| CTAT | 100 |
| ACCA | 100 |
| GTGTG | 100 |
| GGGCG | 99 |
| AACC | 99 |
| TCTCC | 99 |
| AACAA | 99 |
| CAAC | 99 |
| ATAAA | 99 |
| GAAAG | 98 |
| GACC | 98 |
| GCAC | 98 |

| | |
|---|---|
| ACACA | 98 |
| GGGGA | 98 |
| TATC | 97 |
| CCCAC | 97 |
| GAAC | 97 |
| ATAG | 96 |
| GGGCC | 96 |
| GGAAA | 96 |
| AGGGA | 96 |
| CTGGG | 96 |
| GTAA | 95 |
| CCGCC | 95 |
| GTGC | 94 |
| TTTAA | 94 |
| ATAC | 93 |
| AGTA | 93 |
| AAAAC | 93 |
| TAGT | 93 |
| TTCTC | 93 |
| AAGGA | 93 |
| GGGTG | 92 |
| CTGCC | 91 |
| GTTTT | 90 |
| TCCTC | 90 |
| AATAA | 90 |
| AAAAAAA | 90 |
| CCTA | 89 |
| GAGAA | 89 |
| CCACC | 88 |
| AGAAG | 88 |
| CGGGG | 88 |
| GCGGG | 86 |
| CCCAG | 86 |
| TTCCC | 86 |
| GTGGG | 86 |
| GAAGA | 85 |
| TTTCA | 84 |
| TAAC | 84 |
| CTCTG | 84 |
| CCTTT | 84 |
| GGAGA | 83 |
| CCCTG | 82 |
| CTAG | 82 |

| | |
|---|---|
| GAAGG | 81 |
| TTCTG | 80 |
| TTCTTT | 80 |
| GTAT | 79 |
| GGCAG | 79 |
| CTCG | 78 |
| AAAGG | 78 |
| TAAAT | 78 |
| CACAC | 78 |
| TTTCC | 78 |
| GATA | 77 |
| AAGAG | 77 |
| CCAGG | 77 |
| CAGGG | 76 |
| TCTGT | 76 |
| GGTGG | 76 |
| CACCC | 74 |
| GGCTG | 74 |
| CCCTT | 74 |
| CGTG | 73 |
| ATTTA | 73 |
| ACCCC | 73 |
| CGCCC | 73 |
| CAGAG | 73 |
| TTTCTT | 72 |
| TAGG | 71 |
| GGGGT | 71 |
| CGCT | 71 |
| GCAGG | 71 |
| GGCCC | 71 |
| CTAC | 70 |
| CAGCC | 70 |
| GGAT | 69 |
| TGTCT | 69 |
| CCGGG | 69 |
| TTTTCT | 68 |
| AAATT | 68 |
| GGGAA | 68 |
| TCTCA | 68 |
| AAAAAG | 67 |
| AAAGAA | 67 |
| GCCTC | 67 |
| TTTGA | 66 |

| | |
|---|---|
| GGAGGG | 66 |
| ATGC | 66 |
| GCAT | 65 |
| TACC | 65 |
| GGGAGG | 65 |
| AAAGT | 65 |
| CCCTCC | 65 |
| TCTTTT | 65 |
| CACG | 65 |
| CCTGG | 64 |
| TGCCC | 63 |
| TCCG | 63 |
| AAATG | 63 |
| CATTT | 63 |
| AATTT | 62 |
| CCTCCC | 62 |
| TCAAA | 62 |
| CCTGC | 62 |
| ACTA | 62 |
| AAGGG | 62 |
| ATATT | 61 |
| AGGGC | 61 |
| ATTTC | 61 |
| CAGGA | 61 |
| GAAAAA | 61 |
| TAAAAA | 60 |
| TCGG | 60 |
| AAGAAA | 60 |
| AGCG | 60 |
| GAGGC | 59 |
| ACAGA | 59 |
| TCCTG | 59 |
| CGAG | 59 |
| ATATA | 58 |
| CCGT | 58 |
| TCATT | 58 |
| CGGA | 58 |
| TTTTTC | 58 |
| AGAAAA | 57 |
| TTTTTTTT | 57 |
| TGAAA | 56 |
| TATAT | 56 |
| GTCTT | 56 |

| | |
|---|---|
| TGGTT | 55 |
| CGGGC | 55 |
| GGCCG | 55 |
| GTAG | 55 |
| ACGG | 55 |
| GCTGG | 55 |
| CTTTTT | 55 |
| AAAAGA | 54 |
| CGTC | 54 |
| GCCCT | 54 |
| TGAGA | 54 |
| AGCCC | 54 |
| ATTCT | 53 |
| GTCTC | 53 |
| GAAAT | 53 |
| CAAAAA | 53 |
| GTTTG | 53 |
| AATAT | 53 |
| CCCCTC | 53 |
| AGACA | 53 |
| CTCCCC | 52 |
| AGCAG | 52 |
| CCTGT | 52 |
| CCCGG | 52 |
| CCTCA | 52 |
| CTCCA | 52 |
| GACAG | 51 |
| CTGCT | 51 |
| GCCGC | 51 |
| CTGAG | 51 |
| CTGTG | 51 |
| CTTTG | 51 |
| TTCAA | 51 |
| TCGC | 51 |
| TCTGC | 51 |
| CGCA | 51 |
| TTTTTA | 51 |
| TCTTTC | 50 |
| TCCCA | 50 |
| CTCAG | 50 |
| TTGTTT | 50 |
| GGGCGG | 50 |
| AGGCC | 50 |

| | |
|---|---|
| CTGTC | 50 |
| TTCTA | 50 |
| CCAGC | 50 |
| ATTAA | 50 |
| CTCAC | 49 |
| CTCTCT | 49 |
| CTGTT | 49 |
| TGCCT | 49 |
| AGTGA | 49 |
| TCACT | 49 |
| GAGTG | 49 |
| GGGCA | 49 |
| AAAAAT | 49 |
| GGTA | 48 |
| TAATT | 48 |
| CTTTCT | 48 |
| GGCTC | 48 |
| CCGCG | 48 |
| GGCGGG | 48 |
| CCCCGC | 48 |
| CAGAA | 48 |
| TATTA | 48 |
| TGCG | 48 |
| GCAGA | 48 |
| TGGGT | 48 |
| GCCTG | 48 |
| GAGGGG | 48 |
| TGTGG | 47 |
| ACCG | 47 |
| AGAGAG | 47 |
| GGTTT | 47 |
| TTTATT | 47 |
| GTGTGT | 47 |
| CACAG | 46 |
| AATGA | 46 |
| GCCAG | 46 |
| TTAAT | 46 |
| GGGCT | 46 |
| ACTCT | 46 |
| CCGA | 46 |
| TGTGTG | 46 |
| ATCTT | 45 |
| TGCTT | 45 |

| | |
|---|---|
| TGGAG | 45 |
| TCAGA | 45 |
| AAAAAAA | 45 |
| TTCAT | 45 |
| CAGGC | 45 |
| TCTTA | 45 |
| GCCGG | 45 |
| TGGGA | 44 |
| ATTAT | 44 |
| GTGAG | 44 |
| GCGGC | 44 |
| CCACA | 44 |
| GGAGC | 44 |
| CTGGC | 44 |
| GCTCC | 44 |
| GCCCG | 44 |
| TTATTT | 44 |
| ACTTT | 44 |
| TTTGTT | 44 |
| GCGA | 43 |
| AATTA | 43 |
| GTTCT | 43 |
| GGAGGA | 43 |
| GAGGAG | 43 |
| TAGC | 43 |
| AGAAAG | 43 |
| GCGT | 43 |
| CCTCCT | 43 |
| AACAAA | 43 |
| AGTGG | 43 |
| CCCGCC | 43 |
| GGGGCG | 43 |
| TTTGG | 42 |
| TTCTTC | 42 |
| TCTCTC | 42 |
| TCTGG | 42 |
| CCACT | 42 |
| AGGAAG | 42 |
| AGGAGG | 42 |
| CACACA | 42 |
| CTTTA | 42 |
| TAAAG | 42 |
| AATCT | 41 |

| | |
|---|---|
| TTTAC | 41 |
| CTGAA | 41 |
| TTTTTG | 41 |
| GCTTT | 41 |
| CACTC | 41 |
| GCGGGG | 41 |
| AGATG | 41 |
| ACTCC | 41 |
| AAAACA | 41 |
| TTAAAA | 41 |
| AGGGT | 41 |
| TGTTTT | 40 |
| TGAGG | 40 |
| ATGTT | 40 |
| CTGGA | 40 |
| GACG | 40 |
| TGGCC | 40 |
| CAAGA | 40 |
| ATTGT | 40 |
| GTTTC | 40 |
| CTCAT | 40 |
| TTCCTT | 40 |
| CTCCCT | 40 |
| GCTA | 40 |
| TGAGT | 39 |
| GAGCC | 39 |
| AACAG | 39 |
| AAATC | 39 |
| TGGAA | 39 |
| TGTTA | 39 |
| CCGCCC | 39 |
| CGCCCC | 39 |
| ACACAC | 39 |
| ACGC | 39 |
| TTACA | 39 |
| GCCAC | 39 |
| CACTT | 39 |
| CTTCCC | 39 |
| TAATA | 39 |
| GGCCT | 39 |
| AGGCT | 38 |
| TCACC | 38 |
| GCTGC | 38 |

| | |
|---|---|
| AATGT | 38 |
| CTTAA | 38 |
| AGGAAA | 38 |
| ACTTC | 38 |
| AAACAA | 38 |
| TGGCT | 38 |
| AACAT | 38 |
| CTTCCT | 38 |
| TGGGC | 38 |
| ATTTTT | 38 |
| CGCCG | 38 |
| TTTTAA | 38 |
| GCCCA | 38 |
| CAGCT | 38 |
| CGGT | 38 |
| CCTTCC | 38 |
| TCAGT | 38 |
| GAAGT | 37 |
| GTGGT | 37 |
| ACAGG | 37 |
| CCGGC | 37 |
| GAAAC | 37 |
| GAAAGA | 37 |
| GGAAGG | 37 |
| GAGAGA | 37 |
| TCTAT | 37 |
| GAGAC | 37 |
| GGCGC | 37 |
| AGAAT | 37 |
| TTTTTTTT | 37 |
| TCCAT | 37 |
| CCCTCT | 37 |
| CAGTG | 37 |
| TCCCTC | 37 |
| TTTCTTT | 37 |
| AAGCA | 37 |
| TCCAA | 37 |
| ACATT | 37 |
| CCTTG | 36 |
| TGTCC | 36 |
| TCTGA | 36 |
| TTGGT | 36 |
| CCCCCC | 36 |

| | |
|---|---|
| GGCCA | 36 |
| CAAAT | 36 |
| AAATAA | 36 |
| TTCCA | 36 |
| CTTGT | 36 |
| GCGCC | 36 |
| AGTTT | 36 |
| TGTTC | 36 |
| TATTTT | 36 |
| ACCCT | 36 |
| GAGAT | 36 |
| CCCCAC | 36 |
| CAAAG | 36 |
| GCCTT | 36 |
| AGGCA | 36 |
| TATAA | 36 |
| CCAAA | 36 |
| AAAGC | 36 |
| CTCCTT | 36 |
| GGGGGC | 36 |
| TTTTAT | 36 |
| ATTTG | 36 |
| AATTG | 35 |
| CTTCTT | 35 |
| CGCGC | 35 |
| CGCGG | 35 |
| TTCTCT | 35 |
| ACATA | 35 |
| GAGTC | 35 |
| ACCTC | 35 |
| AAAAAAG | 35 |
| TTTGC | 35 |
| CCTCTC | 35 |
| GCAAA | 35 |
| TCTTG | 35 |
| AAAAAC | 35 |
| TTTAAA | 35 |
| TACTT | 35 |
| CGGGGC | 35 |
| TGTAT | 35 |
| TTTAG | 34 |
| GATTT | 34 |
| TTGAA | 34 |

| | |
|---|---|
| CTCTTT | 34 |
| TGTGA | 34 |
| CCCACC | 34 |
| AAAATA | 34 |
| AATAG | 34 |
| CGGCC | 34 |
| CAAAAAA | 34 |
| CAAAC | 34 |
| AGAAGA | 34 |
| TCCTTC | 34 |
| AAGTT | 34 |
| GCCCCC | 34 |
| GTGTT | 34 |
| AAAGAAA | 34 |
| GATC | 34 |
| TCCAC | 34 |
| AAGGAA | 34 |
| AGTCA | 34 |
| AAGAC | 34 |
| GGGGTG | 34 |
| ATGAA | 34 |
| TTGTC | 34 |
| TTATA | 34 |
| CGGGA | 34 |
| CTTAT | 34 |
| AATAAA | 34 |
| GTGGA | 34 |
| CATCT | 33 |
| TCTTCT | 33 |
| AGTGT | 33 |
| CGAC | 33 |
| AGTCT | 33 |
| TAACA | 33 |
| TTAAC | 33 |
| TAAGA | 33 |
| ATACA | 33 |
| TGACA | 33 |
| AAGAT | 33 |
| CTCTCC | 33 |
| GGGGAG | 33 |
| ATAAAA | 33 |
| CCACCC | 33 |
| AGGGAG | 33 |

| | |
|---|---|
| GAGGAA | 33 |
| CTATT | 33 |
| TTTTGT | 33 |
| GTGGGG | 33 |
| GGAGT | 33 |
| TTAAG | 32 |
| TTGTG | 32 |
| AGCCT | 32 |
| AAACT | 32 |
| GAGGGA | 32 |
| GGGAGG[illegible] | 32 |
| CAAGG | 32 |
| CTCAA | 32 |
| GTCCC | 32 |
| ATAAT | 32 |
| TTCTTTT | 32 |
| TTATG | 32 |
| ACCCA | 32 |
| TTTCCT | 32 |
| GAAAAAA | 32 |
| TCAGG | 32 |
| CCCTTC | 32 |
| GGGTC | 32 |
| TGGGGG | 32 |
| ACTCA | 31 |
| TCTCTT | 31 |
| CACCT | 31 |
| ACCTT | 31 |
| AAGGC | 31 |
| ACAAAA | 31 |
| ATGGA | 31 |
| TCATC | 31 |
| TTGCT | 31 |
| TGATT | 31 |
| GTAAA | 31 |
| TGGTG | 31 |
| ATCTC | 31 |
| GCTCT | 31 |
| CCTTCT | 31 |
| TTCG | 31 |
| CACTG | 31 |
| CAGCA | 31 |
| AGCAA | 31 |

| | |
|---|---|
| CAATA | 31 |
| CAGAC | 31 |
| GGGAC | 31 |
| TCCTCC | 31 |
| GTGTGTG | 31 |
| GGTGA | 31 |
| GAAGAA | 31 |
| TTTCTC | 31 |
| TTGAG | 31 |
| AATTC | 30 |
| GTGAA | 30 |
| CCTTA | 30 |
| GCTGA | 30 |
| CCCCCG | 30 |
| ATGAG | 30 |
| CACCA | 30 |
| CTCCTC | 30 |
| ACAAG | 30 |
| CCCAA | 30 |
| TGTTG | 30 |
| ATAGA | 30 |
| TTCAC | 30 |
| TGTGTGT | 30 |
| AAAAAAT | 30 |
| GGGGCC | 30 |
| AGAAC | 30 |
| TGAAG | 29 |
| GTAC | 29 |
| CGGCG | 29 |
| GACTT | 29 |
| AAAAGG | 29 |
| GTTAA | 29 |
| TATGT | 29 |
| TCCAG | 29 |
| CTAAA | 29 |
| AACG | 29 |
| GCAGC | 29 |
| ACTGA | 29 |
| TACAA | 29 |
| CACAA | 29 |
| ACAAT | 29 |
| AGAGAA | 29 |
| CTTCA | 29 |

| | |
|---|---|
| CCTTTT | 29 |
| CCGCT | 29 |
| TATTC | 29 |
| AGAGC | 29 |
| TCCTTT | 29 |
| AGATA | 29 |
| AGAGGA | 29 |
| GGAAAA | 29 |
| GCTTC | 29 |
| AGCCA | 29 |
| CTGCA | 29 |
| TGGCA | 29 |
| CCTCG | 29 |
| TTATC | 28 |
| TATCT | 28 |
| AAGTG | 28 |
| GTTTTT | 28 |
| CATCC | 28 |
| TAAATA | 28 |
| CATTC | 28 |
| TTTTCTT | 28 |
| TCCCCT | 28 |
| ACACACA | 28 |
| AACTG | 28 |
| TGCCA | 28 |
| CCAAC | 28 |
| ATTCA | 28 |
| GTGTC | 28 |
| TTCAG | 28 |
| GGTGT | 28 |
| GAGAGG | 28 |
| AACCT | 28 |
| TTGTA | 28 |
| ACAGT | 28 |
| CTGAC | 28 |
| CCCTCCC | 28 |
| ACGT | 27 |
| ACACT | 27 |
| TGACT | 27 |
| AGAGT | 27 |
| CATAA | 27 |
| TTTCTG | 27 |
| AGAAAAA | 27 |

| | |
|---|---|
| GCCCCG | 27 |
| GAGGT | 27 |
| AGGTG | 27 |
| TGCTG | 27 |
| TGAAT | 27 |
| TCTCCT | 27 |
| GGCAGG | 27 |
| TAGAA | 27 |
| TCTTCC | 27 |
| TGCAG | 27 |
| CCATC | 27 |
| AAAGGA | 27 |
| AAGTA | 27 |
| TCAAT | 27 |
| TGTAA | 27 |
| CAATT | 27 |
| GTATT | 27 |
| TGTTTG | 27 |
| GTTTA | 27 |
| AAAAAAA | 27 |
| GGTGGG | 27 |
| GTCAG | 27 |
| GACAC | 27 |
| GCCCCT | 27 |
| AATGG | 27 |
| CCTGA | 27 |
| AATTTT | 26 |
| TCAGC | 26 |
| CACCCC | 26 |
| GCTGT | 26 |
| GGCGGG( | 26 |
| GCAAG | 26 |
| ACTAA | 26 |
| GACAA | 26 |
| CTCCG | 26 |
| TAGAG | 26 |
| ACAGC | 26 |
| GGGGGG | 26 |
| GGGTT | 26 |
| GACTC | 26 |
| GTCG | 26 |
| AGACT | 26 |
| TATTTA | 26 |

| | |
|---|---|
| GCGCG | 26 |
| TACAT | 26 |
| GGGTGG | 26 |
| TACTC | 25 |
| AGGGGG | 25 |
| ACTGT | 25 |
| TTTTGA | 25 |
| GTCCT | 25 |
| GATGG | 25 |
| TAAAAAA | 25 |
| CTCTA | 25 |
| AAGAGA | 25 |
| AGAGGG | 25 |
| GTCTG | 25 |
| AAAAGAA | 25 |
| TATGA | 25 |
| CCCCCT | 25 |
| GAGCA | 25 |
| TAAAAT | 25 |
| CAACA | 25 |
| GCAGT | 25 |
| TATTG | 25 |
| ATCCC | 25 |
| GAGAAA | 25 |
| TCACA | 25 |
| CTTTTC | 25 |
| AAAGAG | 25 |
| CCCCGCC | 25 |
| TTAAAT | 25 |
| AACCC | 25 |
| TGTCA | 25 |
| TCTCTG | 25 |
| GAATA | 25 |
| TTGGG | 25 |
| TCGT | 25 |
| GGTCT | 25 |
| GGAGGG[illegible] | 25 |
| CGAGG | 25 |
| CCCCCA | 25 |
| CGTGG | 24 |
| CTCTTC | 24 |
| TCTTTCT | 24 |
| ACCAC | 24 |

| | |
|---|---|
| AGTAA | 24 |
| CCAGA | 24 |
| GGGAAG | 24 |
| TAACT | 24 |
| CTTCTC | 24 |
| GACCC | 24 |
| AGGCG | 24 |
| AATCA | 24 |
| TGCCCC | 24 |
| TCTAA | 24 |
| ACGA | 24 |
| CACACAC | 24 |
| AGGTT | 24 |
| GAAGC | 24 |
| ATGGG | 24 |
| TTTTTCT | 24 |
| GGACC | 24 |
| CCCGCCC | 24 |
| CATTA | 24 |
| AACAC | 24 |
| AAGCC | 24 |
| TTCCCT | 24 |
| TTAGA | 24 |
| TTTTTTA | 24 |
| GTGGC | 24 |
| GGGGGT | 23 |
| ATTTAT | 23 |
| AAAATT | 23 |
| TTGAT | 23 |
| ATATAT | 23 |
| GTCAC | 23 |
| ATCCT | 23 |
| AGCTG | 23 |
| AGGAC | 23 |
| TAGTT | 23 |
| AAGAAG | 23 |
| CTCGG | 23 |
| CACAT | 23 |
| GTTCC | 23 |
| CCGCCCC | 23 |
| AAGAAAA | 23 |
| CTTGG | 23 |
| GGAAC | 23 |

| | |
|---|---|
| GAAGGA | 23 |
| GCACC | 23 |
| CCTCTG | 23 |
| CCCCTCC | 23 |
| CAGCCC | 23 |
| GAATG | 23 |
| CGAA | 23 |
| GGGGGA | 23 |
| GAGAAG | 23 |
| AAACAAA | 23 |
| CCACG | 23 |
| TCCTA | 23 |
| TATCA | 23 |
| GCAGGG | 23 |
| GGGAGA | 23 |
| GATGA | 23 |
| ATATTT | 23 |
| ATTAAA | 23 |
| ATGTA | 23 |
| CTCTGT | 23 |
| GGGCTG | 23 |
| CCCAGC | 23 |
| ACTGC | 23 |
| GGTCC | 23 |
| CCCCAG | 23 |
| GAGCT | 23 |
| CCATG | 23 |
| CTGCCC | 23 |
| TTACT | 23 |
| ACCCCC | 23 |
| TTTATTT | 23 |
| TGTGTGT | 23 |
| GTTGT | 22 |
| CCTCTT | 22 |
| TCCCG | 22 |
| TCCGC | 22 |
| CGCTC | 22 |
| GGGCGG | 22 |
| CAAAAAA | 22 |
| TCATA | 22 |
| CCCAT | 22 |
| CCAGGC | 22 |
| GCTGGG | 22 |

| | |
|---|---|
| TGCTC | 22 |
| GACCT | 22 |
| CACGG | 22 |
| CAGGT | 22 |
| TCTTTTT | 22 |
| CAGGGA | 22 |
| GGGGCG( | 22 |
| GAACA | 22 |
| TTTGTTT | 22 |
| TTGTTTT | 22 |
| GGCTGG | 22 |
| AGGGGA | 22 |
| AGATT | 22 |
| GCCGGG | 22 |
| TCAAG | 22 |
| CCAAG | 22 |
| GAATT | 22 |
| TTCTTTC | 22 |
| GTGAC | 22 |
| CCCAGG | 22 |
| GAAAAG | 22 |
| CAGTT | 22 |
| TCCTCT | 21 |
| CGTT | 21 |
| ATGTG | 21 |
| TCAAAA | 21 |
| TCCCCC | 21 |
| GAGAGA( | 21 |
| GCCTCC | 21 |
| GGAAT | 21 |
| GCAGGA | 21 |
| ATATG | 21 |
| CGGGGG | 21 |
| AAAAAAA | 21 |
| GTCTTT | 21 |
| ATTAC | 21 |
| CCCGCG | 21 |
| CAGAT | 21 |
| CCGTG | 21 |
| ATAGT | 21 |
| CATGT | 21 |
| GATAA | 21 |
| ATCAA | 21 |

| | |
|---|---|
| CAACC | 21 |
| TTCCTC | 21 |
| ATGAT | 21 |
| CCATT | 21 |
| GGACT | 21 |
| AGGGAA | 21 |
| GGACA | 21 |
| CTCTCTC | 21 |
| AGTTC | 21 |
| CCGGGC | 21 |
| CCTCCCC | 21 |
| AGTTA | 21 |
| GTGTGTG | 21 |
| CGGAG | 21 |
| CTCGC | 21 |
| GGGGAA | 21 |
| GTAAT | 20 |
| CAGTC | 20 |
| TCATG | 20 |
| AGAGAGA | 20 |
| ATAAC | 20 |
| AAAAGT | 20 |
| ATCAG | 20 |
| ATAAG | 20 |
| TTGGC | 20 |
| TTTTTTG | 20 |
| ATCCA | 20 |
| TGGTTT | 20 |
| ATCAC | 20 |
| TACAG | 20 |
| ACACC | 20 |
| ATGCC | 20 |
| AAGTC | 20 |
| GTCAT | 20 |
| TTTTCA | 20 |
| TAATG | 20 |
| CCGGGG | 20 |
| TTTCAA | 20 |
| TCGGG | 20 |
| GGAGAA | 20 |
| CATTTT | 20 |
| CCCTCTC | 20 |
| TTTTTTC | 20 |

| | |
|---|---|
| GGAGAG | 20 |
| GTTAT | 20 |
| ATTCTT | 20 |
| AAATGA | 20 |
| GTTGG | 20 |
| CTTTTTT | 20 |
| TCTCCC | 20 |
| TTTTATT | 20 |
| TGAGC | 20 |
| AGGGTG | 20 |
| GGCAC | 20 |
| GGGCCC | 20 |
| TTTTTTTT | 20 |
| GGTTC | 20 |
| AAGAGG | 20 |
| ATACT | 19 |
| GGATT | 19 |
| TTGCA | 19 |
| AAAAAGA | 19 |
| TATATA | 19 |
| CTCCCA | 19 |
| ATCTG | 19 |
| CATAT | 19 |
| AAGGT | 19 |
| CAATG | 19 |
| TGACC | 19 |
| GGAGGA( | 19 |
| TAAAC | 19 |
| TTCTGT | 19 |
| ATTTCT | 19 |
| GACAT | 19 |
| ACAGAA | 19 |
| CTGGGG | 19 |
| GCGGA | 19 |
| TGTCTT | 19 |
| GGCTT | 19 |
| GCGTG | 19 |
| TGTGC | 19 |
| CTGGGC | 19 |
| CAGGAG | 19 |
| CCTCCTC | 19 |
| ATTCC | 19 |
| GGCAA | 19 |

| | |
|---|---|
| CCAGGG | 19 |
| CACCG | 19 |
| TTAGT | 19 |
| TTGGTT | 19 |
| TTCTTCT | 19 |
| TTCCCC | 19 |
| CTTAG | 19 |
| AGCTT | 19 |
| GTTTGT | 19 |
| TCTAG | 19 |
| TCTCTCT | 19 |
| CTAAT | 19 |
| ATTTAA | 19 |
| TTTTCC | 19 |
| CGCAG | 19 |
| TCCCCA | 19 |
| TCCCTT | 19 |
| AATAC | 19 |
| GGGCCG | 19 |
| ATTGA | 19 |
| GCGGGG( | 19 |
| TTTTTAA | 19 |
| TGCAT | 18 |
| ATTTTA | 18 |
| CTTGA | 18 |
| TGAAAA | 18 |
| CCCGT | 18 |
| TTTGGT | 18 |
| CTGTA | 18 |
| GTGCC | 18 |
| CCGCGC | 18 |
| ACTTA | 18 |
| TCTAC | 18 |
| CTAAG | 18 |
| GGAGGC | 18 |
| CTGCCT | 18 |
| AGAAAGA | 18 |
| AACTC | 18 |
| CAGAAA | 18 |
| AGCAC | 18 |
| TGGGAG | 18 |
| CACACAC/ | 18 |
| ATAAAT | 18 |

| | |
|---|---|
| CTGGCC | 18 |
| ATATC | 18 |
| TCAAC | 18 |
| CTTTAT | 18 |
| CCGAG | 18 |
| CTTTCTT | 18 |
| ACGGG | 18 |
| TGCAA | 18 |
| CCCCTT | 18 |
| AAAAACA | 18 |
| CTGAT | 18 |
| GCGAG | 18 |
| TTAAAAA | 18 |
| CGCTG | 18 |
| TATTAT | 18 |
| GGCCGG | 18 |
| GAAGGG | 18 |
| CCAGCC | 18 |
| CGCAC | 18 |
| CCCTGC | 18 |
| CCTGCC | 18 |
| CGGGAG | 18 |
| TGTGTGT | 18 |
| TTACC | 17 |
| TGATG | 17 |
| CCTAC | 17 |
| CCGTC | 17 |
| AGGCTG | 17 |
| GGCGGC | 17 |
| AAACC | 17 |
| AACTT | 17 |
| TGCCTC | 17 |
| ATTTTC | 17 |
| AAGAAA | 17 |
| CCTGGG | 17 |
| ACACACA | 17 |
| TCTTTG | 17 |
| CCCCGG | 17 |
| GGGAAA | 17 |
| GCCGCC | 17 |
| CAATC | 17 |
| TTCAAA | 17 |
| CCAGT | 17 |

| | |
|---|---|
| TCATTT | 17 |
| CATCA | 17 |
| CCACCCC | 17 |
| AGTCC | 17 |
| GACTG | 17 |
| AAGGAG | 17 |
| ATGTTT | 17 |
| TAGAT | 17 |
| CCTAG | 17 |
| TTTTGTT | 17 |
| AAAAAAC | 17 |
| CGGGGCC | 17 |
| TTATTA | 17 |
| TCTGCC | 17 |
| CCCGA | 17 |
| GGTTG | 17 |
| CAAGT | 17 |
| GAGGGC | 17 |
| CTGGT | 17 |
| ATTATT | 17 |
| TTTGAG | 17 |
| ACCTG | 17 |
| TTTCCTT | 17 |
| GGTGC | 17 |
| CTTTCC | 17 |
| CGCCGC | 17 |
| GTTCTT | 17 |
| AAAAAAA | 17 |
| GTGTGTG | 17 |
| GATTA | 16 |
| GTTTTC | 16 |
| TGTTTTT | 16 |
| TACCT | 16 |
| GATGT | 16 |
| CTTCTG | 16 |
| GCTCA | 16 |
| AAATTA | 16 |
| CCTAT | 16 |
| GAAGAG | 16 |
| ATGGT | 16 |
| AGAGAGA | 16 |
| TAAGT | 16 |
| TGGAGG | 16 |

| | |
|---|---|
| CAGGAA | 16 |
| AAATAT | 16 |
| CTCCCCT | 16 |
| ACACACA | 16 |
| GTCAA | 16 |
| AACTA | 16 |
| GCCAA | 16 |
| GGAAGA | 16 |
| TCGCC | 16 |
| AGGAT | 16 |
| GGCCCC | 16 |
| TTGAC | 16 |
| AGTAG | 16 |
| ACATG | 16 |
| CCCCGCC | 16 |
| ACCAT | 16 |
| TATAAA | 16 |
| GATTC | 16 |
| GTCTCT | 16 |
| CTTAC | 16 |
| CCAAT | 16 |
| CTGCTT | 16 |
| AGAAGA | 16 |
| CAGACA | 16 |
| GAGGGA | 16 |
| CCCGGG | 16 |
| TGGGGA | 16 |
| ACTTG | 16 |
| AGGTC | 16 |
| CTCCCTC | 16 |
| CCTAA | 16 |
| TTCTCC | 16 |
| CGGTG | 16 |
| CTTTTA | 16 |
| TCTGTG | 16 |
| CCCGGC | 16 |
| ACCGC | 16 |
| GTGCT | 16 |
| TCCCTG | 16 |
| TGGTC | 16 |
| CCTTTC | 16 |
| CCCTCA | 16 |
| GAACC | 16 |

| | |
|---|---|
| AACAAAA | 16 |
| AAGGGG | 16 |
| AGGGGC | 16 |
| GGGAT | 15 |
| GAGTT | 15 |
| CTTTTG | 15 |
| AAATTT | 15 |
| CAGGGG | 15 |
| GCGGGC | 15 |
| CGGGCG | 15 |
| CTACC | 15 |
| GGCCGC | 15 |
| GCCGCG | 15 |
| AACCA | 15 |
| GGGCCA | 15 |
| TTAATT | 15 |
| ACCAG | 15 |
| AAGAAGA | 15 |
| GGTAA | 15 |
| AAAGTT | 15 |
| GTAAG | 15 |
| CTTGC | 15 |
| GGGCGG( | 15 |
| AAAATG | 15 |
| TATATT | 15 |
| AAAATC | 15 |
| CTGCG | 15 |
| GCCCTG | 15 |
| CGTCC | 15 |
| GAAACA | 15 |
| CTCACC | 15 |
| AGGAGA | 15 |
| CAACT | 15 |
| GGGCAG | 15 |
| GAGTGA | 15 |
| AGTGAA | 15 |
| CTTTTCT | 15 |
| TCCCTCC | 15 |
| TCTTTA | 15 |
| TCCGG | 15 |
| CCGCCG | 15 |
| TTTCTTTT | 15 |
| CTATA | 15 |

| | |
|---|---|
| GGATG | 15 |
| GCCAGG | 15 |
| CAGGCC | 15 |
| CTAGA | 15 |
| GGGGCT | 15 |
| TGTTTC | 15 |
| GCCTGG | 15 |
| GCCCAG | 15 |
| TTTGAA | 15 |
| TGATA | 15 |
| TCTTCA | 15 |
| TAGCA | 15 |
| CGGCA | 15 |
| CAGCG | 15 |
| CTACA | 15 |
| ACTTTT | 15 |
| AGACC | 15 |
| CAGCCT | 15 |
| CCTGGC | 15 |
| GTGGGG[illegible] | 15 |
| GACCA | 15 |
| GGGGAG[illegible] | 15 |
| GCACT | 15 |
| GTTGA | 15 |
| CTCCTG | 15 |
| ACAAAAA | 15 |
| TATTTTT | 15 |
| ATAAAAA | 15 |
| TCCTTCC | 15 |
| AGTGC | 15 |
| GAGCG | 15 |
| GCACA | 15 |
| TTTCTCT | 15 |
| CTTCTTT | 15 |
| GAGGAG[illegible] | 15 |
| GGGCGC | 15 |
| AGAAGG | 15 |
| CCCCCAC | 15 |
| CCTTCCC | 15 |
| TGTGTT | 15 |
| GTTCA | 15 |
| GGTCA | 15 |
| TGTTCT | 15 |

| | |
|---|---|
| GCCCCGC | 15 |
| TGTGTGT( | 15 |
| GAAAGG | 14 |
| ATGTC | 14 |
| AGTAT | 14 |
| GTTAC | 14 |
| TCCCAC | 14 |
| AAATAG | 14 |
| CGGCGG | 14 |
| TACCC | 14 |
| CATGG | 14 |
| AGGAGC | 14 |
| CTTAAA | 14 |
| CGCCCCC | 14 |
| AAAGAAA | 14 |
| TATCC | 14 |
| CCCCTG | 14 |
| GCTCCC | 14 |
| AGGGCG | 14 |
| ATATTA | 14 |
| TTCCTG | 14 |
| TCCCCG | 14 |
| CTAGG | 14 |
| CATTG | 14 |
| ACATC | 14 |
| AGGAAGC | 14 |
| GTAGA | 14 |
| TCTCAT | 14 |
| ATTGG | 14 |
| CAGAGG | 14 |
| CCTCAG | 14 |
| TTTCAT | 14 |
| AAAACAA | 14 |
| TCTTAA | 14 |
| GATAT | 14 |
| TTCATT | 14 |
| GGAAAT | 14 |
| TACTG | 14 |
| AATATA | 14 |
| TGGAT | 14 |
| AGCCG | 14 |
| ATACC | 14 |
| ACAAC | 14 |

| | |
|---|---|
| CACTA | 14 |
| GTCCA | 14 |
| ATGCT | 14 |
| GAAAAAA | 14 |
| TGCTTT | 14 |
| TTTAAT | 14 |
| AGAGAGG | 14 |
| TGGGCC | 14 |
| CTCTGC | 14 |
| TTGCC | 14 |
| AATATT | 14 |
| ATCG | 14 |
| CGCCT | 14 |
| AGCCCC | 14 |
| AAACAT | 14 |
| AAAAAAA | 14 |
| GCCCGC | 14 |
| TATAC | 14 |
| CTGGGT | 14 |
| CTCCTTC | 14 |
| CCTCCCT | 14 |
| TTTTAAA | 14 |
| TCCTTTT | 14 |
| GGCGGGG | 14 |
| TAAAAG | 14 |
| ACGCC | 14 |
| CACGC | 14 |
| CGTCT | 14 |
| TAGGT | 13 |
| AGTTG | 13 |
| TGTCTC | 13 |
| GAAAAT | 13 |
| ACCCG | 13 |
| GGCTGC | 13 |
| TGTGAG | 13 |
| GATTG | 13 |
| TCCTTA | 13 |
| GTTGC | 13 |
| CCGCGG | 13 |
| TAGGA | 13 |
| CAAAAAA | 13 |
| CTCCCCC | 13 |
| TAATTT | 13 |

| | |
|---|---|
| GGAAAG | 13 |
| GAAAGAA | 13 |
| GCTCG | 13 |
| AGAGAGA | 13 |
| GAGAGAG | 13 |
| AGGTA | 13 |
| TTAACT | 13 |
| ACTTCC | 13 |
| TCAGAG | 13 |
| TAAACA | 13 |
| GACGG | 13 |
| CGGCT | 13 |
| TCCCCTC | 13 |
| GAGATG | 13 |
| ACCAA | 13 |
| CTACT | 13 |
| GGCGT | 13 |
| CAGAGA | 13 |
| CAAACA | 13 |
| CTCTCCT | 13 |
| GCCGT | 13 |
| CGGGCC | 13 |
| CGGCCC | 13 |
| CTCCTCC | 13 |
| CATTTC | 13 |
| ATCTTT | 13 |
| GCCCCA | 13 |
| AGCAGA | 13 |
| TTCTTA | 13 |
| ACAAAG | 13 |
| CGGGT | 13 |
| AAACAC | 13 |
| GAAATG | 13 |
| AAGAGAA | 13 |
| AAGGAAA | 13 |
| GTGCA | 13 |
| TAGTG | 13 |
| TTTGGTT | 13 |
| CTCCAA | 13 |
| CCCTA | 13 |
| TGCGG | 13 |
| CTGGGA | 13 |
| CTTCCCT | 13 |

| | |
|---|---|
| CCTTCTT | 13 |
| CATGA | 13 |
| CATGC | 13 |
| ATGAC | 13 |
| AAGGGA | 13 |
| TATAG | 13 |
| ACTGG | 13 |
| CCTGCT | 13 |
| TCTGTC | 13 |
| GGCCTC | 13 |
| TTTAAAA | 13 |
| ACAGAG | 13 |
| GGACG | 13 |
| AAAATAA | 13 |
| TGAGGG | 13 |
| TTGTCT | 13 |
| TTAACA | 13 |
| TAATAA | 13 |
| CCTCTCT | 13 |
| TATTTG | 13 |
| GTAGG | 13 |
| GGCGCG | 13 |
| AAGCT | 13 |
| GCCCGG | 13 |
| ATTTTTT | 13 |
| CCCACT | 13 |
| CCTTCCT | 13 |
| GGGGGTG | 13 |
| GACACA | 13 |
| TGGCG | 13 |
| TGAGAA | 13 |
| CCCTTT | 13 |
| GTTTTG | 13 |
| TTATTTT | 13 |
| GCCCCTC | 13 |
| TTTTTAT | 13 |
| GTGTGTG | 13 |
| GGGGGCC | 13 |
| CCCTCCC | 13 |
| TCGA | 13 |
| CTCTTTC | 12 |
| GGCGA | 12 |
| CTCAGC | 12 |

| | |
|---|---|
| TAAAAAA | 12 |
| TGCCCT | 12 |
| GCGCT | 12 |
| GCGGCC | 12 |
| ATAGG | 12 |
| CCCCCGC | 12 |
| GAGTA | 12 |
| TGTTAA | 12 |
| TCCTTCT | 12 |
| ACACAT | 12 |
| GCGCCC | 12 |
| GGGAGGA | 12 |
| AAGTTT | 12 |
| CTGCCTC | 12 |
| GAATC | 12 |
| TATTTC | 12 |
| AAGAAAA | 12 |
| TTTTTTTG | 12 |
| CATAC | 12 |
| GAAATA | 12 |
| CCCTGG | 12 |
| TGGGGC | 12 |
| GGAAGGA | 12 |
| AAGGAAG | 12 |
| CACCCT | 12 |
| CTTCTA | 12 |
| TCCTGT | 12 |
| AAAAAAA | 12 |
| AAAAAGA | 12 |
| CTCAAA | 12 |
| CCCCACC | 12 |
| AAGACA | 12 |
| CCTCTCC | 12 |
| CTGCCA | 12 |
| AATGAA | 12 |
| GGGGCA | 12 |
| AGCGC | 12 |
| GACAAA | 12 |
| GGGTGT | 12 |
| AGAGGC | 12 |
| TAGGG | 12 |
| CCCGCCC | 12 |
| CCCACCC | 12 |

| | |
|---|---|
| GCAGAG | 12 |
| GAAAAC | 12 |
| TTAAAG | 12 |
| TGAAC | 12 |
| AAAGCA | 12 |
| AAGCAG | 12 |
| AGGAAAA | 12 |
| GGAAAAA | 12 |
| AAAGTA | 12 |
| GTATC | 12 |
| CCATA | 12 |
| TCTGTT | 12 |
| GCATG | 12 |
| TTTTTGT | 12 |
| CACTCC | 12 |
| CTGTTT | 12 |
| TTTCAC | 12 |
| AAAAGC | 12 |
| AATTAA | 12 |
| CTTTGT | 12 |
| TTTGTA | 12 |
| TTGGA | 12 |
| TTGTTG | 12 |
| AACATT | 12 |
| TGTTTA | 12 |
| AGCAAA | 12 |
| AAAAGGA | 12 |
| AGCTC | 12 |
| TCCTGC | 12 |
| GTGGGA | 12 |
| GCCTA | 12 |
| GGTGGG | 12 |
| ATTTGT | 12 |
| GGCAGG | 12 |
| GCTTG | 12 |
| AATAAT | 12 |
| CCTGTG | 12 |
| AAATCT | 12 |
| GGTTTT | 12 |
| CACACAC | 12 |
| AAATGG | 12 |
| AGTGTG | 12 |
| GGCAT | 12 |

| | |
|---|---|
| TGTTTGT | 12 |
| GCATT | 12 |
| TTACTC | 12 |
| GCAAT | 12 |
| TCTTTTC | 12 |
| TGGGTG | 12 |
| AAAAAGG | 12 |
| CAGCAG | 12 |
| CAGGGC | 12 |
| GGAGTG | 12 |
| GCTAG | 12 |
| TTTCTTTC | 12 |
| CCCTTCC | 12 |
| AAAGGAA | 12 |
| GTCGG | 12 |
| CTGTCT | 12 |
| GCGGT | 12 |
| TTGTTA | 12 |
| TTCTTTCT | 12 |
| CTTTCTC | 12 |
| TTCTCA | 12 |
| ACTTCT | 12 |
| TTTTTTAA | 12 |
| AGGCAG | 12 |
| CTGCTG | 12 |
| TGAGAG | 12 |
| AAAAATA | 12 |
| AAATAAA | 12 |
| CACAGA | 12 |
| CTAAC | 12 |
| GAACT | 12 |
| GTGCG | 12 |
| ATCAT | 12 |
| TTTGAT | 12 |
| TAACAA | 12 |
| TGAAGA | 12 |
| GAAAAGA | 12 |
| TGTGGG | 12 |
| CGCTCC | 12 |
| AAAAAAA | 12 |
| CCACCT | 12 |
| TGTGTGT | 12 |
| GAGGGG | 12 |

| | |
|---|---|
| AGAGAGA | 12 |
| GAGAGAG | 12 |
| TTAGG | 11 |
| TCTTCTT | 11 |
| TTTATC | 11 |
| GTGGTT | 11 |
| AATTCT | 11 |
| TGGTA | 11 |
| CAAAAAA | 11 |
| GGCCAG | 11 |
| TGGAAA | 11 |
| CTATG | 11 |
| CTGAGA | 11 |
| TGAGAC | 11 |
| CTGGAG | 11 |
| TTTCG | 11 |
| AAAAGAA | 11 |
| ACAGAAA | 11 |
| AGAAAT | 11 |
| AGGAGGA | 11 |
| CTCTGG | 11 |
| GCAAGA | 11 |
| AACATA | 11 |
| ACATAA | 11 |
| GCAAGG | 11 |
| ACAATA | 11 |
| TCTCAA | 11 |
| TTTGCT | 11 |
| GGCAGA | 11 |
| TTTTGC | 11 |
| GTCATT | 11 |
| CAGGCT | 11 |
| TGGGGT | 11 |
| TAAGG | 11 |
| AGCGA | 11 |
| CTTTCA | 11 |
| CCCCTCT | 11 |
| TTTTTTTC | 11 |
| CTCATT | 11 |
| TATCTT | 11 |
| ACAAAT | 11 |
| GCCGA | 11 |
| ACCCCT | 11 |

| | |
|---|---|
| TAATC | 11 |
| AATCTT | 11 |
| TCATCT | 11 |
| ATCTA | 11 |
| CTTCCTT | 11 |
| GCTTA | 11 |
| GAGGCA | 11 |
| ATGAAA | 11 |
| GTTTTTT | 11 |
| AAAAAGT | 11 |
| TCTATT | 11 |
| TGTAG | 11 |
| TTAAGA | 11 |
| TAAGAA | 11 |
| GTCTA | 11 |
| CCCTTCT | 11 |
| GGTTA | 11 |
| TTTGTG | 11 |
| GACTTT | 11 |
| CTCTGA | 11 |
| TGTATT | 11 |
| ATTGTT | 11 |
| TTGTTTG | 11 |
| ATTTCA | 11 |
| TCTCTTT | 11 |
| AAATGT | 11 |
| GTTAAA | 11 |
| AGAAAAA | 11 |
| TTTATA | 11 |
| GAAGAAG | 11 |
| TGATC | 11 |
| GATTTT | 11 |
| CCTGTC | 11 |
| TGGAC | 11 |
| ACCCCA | 11 |
| GGCCCG | 11 |
| ATTAG | 11 |
| AGCAT | 11 |
| GCATA | 11 |
| CACCAC | 11 |
| GACAGA | 11 |
| TTCTGC | 11 |
| GTTATT | 11 |

| | |
|---|---|
| GTGTTT | 11 |
| TTATAT | 11 |
| TTAAATA | 11 |
| ACACACA | 11 |
| CACACAC | 11 |
| CCTCAC | 11 |
| AAATTC | 11 |
| CAGTGT | 11 |
| TAAAGA | 11 |
| TTTTCTTT | 11 |
| AGGCCC | 11 |
| AGCGG | 11 |
| GCGCGG | 11 |
| GTGTA | 11 |
| AAGCAA | 11 |
| CCTTTTT | 11 |
| CTGTGT | 11 |
| TATTATT | 11 |
| TTTTGAG | 11 |
| TATGC | 11 |
| ATGCA | 11 |
| CTTCCTC | 11 |
| TCCTCCC | 11 |
| CACTTC | 11 |
| GAGGAA | 11 |
| GCCACC | 11 |
| CTTTCTTT | 11 |
| GTAAAA | 11 |
| GGAAGT | 11 |
| TTTAAC | 11 |
| AGAGGA | 11 |
| AGAGTG | 11 |
| TTCTAA | 11 |
| CTAAAA | 11 |
| CTAGT | 11 |
| GGAGCC | 11 |
| CCGGA | 11 |
| ATTTAC | 11 |
| AGACAG | 11 |
| AGGCGG | 11 |
| AACAGA | 11 |
| CTCCAC | 11 |
| TTTCTTC | 11 |

| | |
|---|---|
| GAGAAAG | 11 |
| GGGTGGG | 11 |
| TAGTC | 11 |
| GGGGCGG | 11 |
| GCGGGGG | 11 |
| AAAGGG | 11 |
| GTGTGTG | 11 |
| TCTGGG | 11 |
| AGAGAGA | 11 |
| AGAGAGA | 11 |
| GAGAGAG | 11 |
| GAGAGAG | 11 |
